# A three-strain synthetic community model whose rapid response to antagonism allows the study of higher-order dynamics and emergent properties in minutes

**DOI:** 10.1101/2022.09.07.505616

**Authors:** Bernardo Aguilar-Salinas, Gabriela Olmedo-Álvarez

**Affiliations:** Departamento de Ingeniería Genética, Unidad Irapuato, Centro de Investigación y Estudios Avanzados del IPN, Irapuato, México

**Keywords:** higher-order interactions_1_, bacterial antagonism_2_, synthetic community_3_, emergent properties_4_, immediate response_5_

## Abstract

A fundamental question in the assembly of microbial communities is how complex systems arise from a few components. Synthetic communities allow addressing the dynamics and mechanisms of complex microbial interactions. Most studies on microbial interactions are done in lapses of hours and even days, but microbes must be able to sense danger in seconds. We assembled a three-strain synthetic community from the phylum Bacillota that, based on previously evaluated paired interactions, appear to have different ecological roles: resistant (R), antagonists (A), and sensitive (S). The BARS synthetic community (*Bacillota* A + S + R) reproduces features of complex communities and exhibits higher-order interaction dynamics. The majority of the S strain population, *Sutclifiella horikoshii* 20a, dies within 5 min in a paired interaction with A strain, *Bacillus pumilus* 145. An emergent property appears upon adding the third interactor, as antagonism of strain A over S is not observed in the presence of the R strain, *Bacillus cereus* 111. After the first five min a change of state of the cells is observed, as the surviving population of the S strain seemed to have acquired tolerance to A. In summary, our model allows the study of the assembly dynamics of a three-species community and to evaluate the immediate outcome within a 30 min frame. The BARS has features of a complex system where the paired interactions do not predict the community dynamics. The model is amenable to mechanistic dissection and to modeling how the parts integrate to achieve collective properties.

**IMPORTANCE:** Microbial communities are of utmost importance, given their roles in health, agriculture, and all biogeochemical cycles on Earth. Synthetic ecology studies communities by reducing the number of variables. Microbial interactions are usually evaluated in hours or days, however, upon a first encounter, bacteria must respond in minutes, particularly when competition involves killing of neighboring cells. We generated a synthetic community of three species that allows the study of community dynamics in a 30 min frame. We denominated our model BARS as it comprises Bacillota strains that in paired interactions are Antagonist, Resistant, or Sensitive. Even though in paired interaction the antagonist kills the sensitive strain, in a triple interaction, the resistant strain provides stability to the community by neutralizing the antagonism. Therefore, BARS is a rapid response model with features of a complex system where the paired interactions do not predict the community dynamics and exhibit emergent properties.

## INTRODUCTION

Microbial communities form the basis of all biological systems and are dynamic and complex. The complexity of microbial communities results, among other things, from the great diversity of interacting species. Studying bacterial communities in nature represents a significant challenge, given the numerous biotic and abiotic variables that need to be considered. Extensive effort has been dedicated to understanding the processes that contribute to community assembly (Reviewed in Nemergut *et al*., 2013). Synthetic ecology offers an approach to studying communities, reducing the number of variables, particularly the number of interacting species, the environment, and the spatial setting. One critical aspect, though often underappreciated, is that synthetic ecology allows controlling the timing of the experiments.

Different individuals in a community have distinct genetic repertoires that can be deployed as specific phenotypes to cover immediate needs (space and resources) and, more importantly, to react to perceived threats, which is critical for survival. In communities, competitive interactions are unavoidable (Hibbing *et al*., 2010; Cornforth & Foster, 2013). For example, the efficient use of the nutrients by an individual may leave another without nutrients; for the acquisition of iron, individuals that produce siderophores are at an advantage for iron-scavenging (Wandersman & Delepelaire, 2004). On the other hand, direct competition occurs when a species inhibits the growth of another by producing extracellular molecules (interference competition). This type of competition has been amply studied, particularly as it involves the synthesis of antibiotics and surfactants (Cornforth & Foster, 2013; Niehus *et al*. 2021). A common approach to studying interactions among microbes is through the analysis of pairwise interactions. This strategy was, after all, the source of discovery of natural products encoded by bacteria and fungi that have been the sources of many antibiotics in current clinical use. Antimicrobial compounds are the best and most studied example of interference competition, as these molecules can mediate interactions either between different species (Stubbendieck & Straight, 2015), between related strains of the same species (Lyons & Kolter, 2017) or between genetically identical individuals in a population (Kerr *et al*., 2002). Bacteria can have systems for delivering toxins to neighboring bacteria upon direct cell contact. Well-known systems are type VI secretion (T6S) and contact-dependent growth inhibition (CDI) systems (Garcia, E.C., 2018). If bacteria have evolved such variable ammunition for competition, how can community diversity be maintained?

Modeling pairwise species interactions has been a starting point in many works to model the complexity and dynamics of multispecies communities. Carrara *et al*. (2015) simulated data derived from an empirical interaction matrix to predict multispecies community dynamics across multiple species (ten protists and one rotifer). They found some limitations in inferring from paired interactions when applied to organisms with longer generation times, mainly since the data depended on the experimental sampling times (Carrara *et al*., 2015). Friedman *et al*. 2017 evaluated paired interactions and then generated communities with three or more members. They suggest that a rule can be defined from the results observed in pairs and that in a multispecies setting, species that all coexist with each other in pairs will survive, while species that are excluded by any surviving species will go extinct (Friedman *et al*., 2017). An important observation that makes prediction from paired interactions complicated is that interactions may be modulated by the presence of additional species (Billick, I. & Case, T. J. 1994).

Different simplified microbial models have been used to explain the coexistence and assembly of communities. A classic example of a simplified community is the nontransitive rock-paper-scissors model by Kerr (2002), which is based on a community of three *E. coli* isogenic strains (toxin-producer, toxin-sensitive, and toxin-resistant strain). In this work, it was shown that coexistence occurs where there is structure (i.e. agar plates), where patches of the different strains can form, but not in a mixed environment (i.e., laboratory flask) (Kerr *et al*., 2002). Another synthetic community is the “THOR” model by

Lozano *et al*. (2019), a three-strain community in which two of the interactors have an antagonistic interaction, but, with the addition of a third, this interaction is modified, and antagonism does not occur (Lozano *et al*., 2019). In this model, the responses are evaluated at ten hours and up to two days.

The principal feature of the models mentioned above is that each comprises three organisms. Although this may not seem very complex, these models show emergent properties that are not observed in paired interactions but emerge in communities of three or more species. Some models represent non-transitive interactions, in which the interactions could occur indirectly and be non-hierarchical, as is observed in Higher Order Interactions (HOIs). HOIs arise with the increased complexity in bacterial communities. Three or more organisms must be present for high-order interactions to emerge, and in these interactions, the effect of one competitor on another depends on the presence of a third species.

The term “higher-order interactions” has been defined in various ways in the ecological literature (Billick, I. and Case, T.J., 1994). One use of the term HOI implies that indirect interactions arise, for instance, when the impact of one species on another requires or is modified by the presence of a third species (Wootton 1994). Studying the species in isolation or in pairs cannot predict many functional changes. The term HOI is also employed if a paired interaction fails to predict the dynamics of the community. HOIs have to be considered an intrinsic property of complex microbial communities. The study of HOI is essential to understand the dynamics of the communities and what allows species coexistence and makes them robust to the interactions between them and the environment (Grilli *et al*. 2017). In a higher-order interaction, one competitor modifies the competition between the other two (Levine *et al*. 2017). The occurrence of HOIs in bacterial communities has been observed, although sometimes it was not described as an HOI (Gallardo-Navarro & Santillan, 2019; Lozano *et al*., 2019; Piccardi *et al*., 2019; Mickalide & Kuehn, 2019; Sanchez-Gorostiaga *et al*., 2019).

Communities are continually perturbed in various manners, with dispersion and invasion forcing new neighbors and the need to respond to potentially harmful interactors. Most studies on microbial interactions are done in lapses of hours or even days (Vaz & Kinkel, 2014; Kerr *et al*., 2002; Kirkup & Riley, 2004). The time of a response is an essential factor in competence, and a delay in responding, to antagonism, for instance, could lead to the death of a population. In the works cited above, responses to the interactions were measured in long-term experiments that, in some cases, ranged between 10 and 50 hours (Lozano *et al*., 2019), and in other cases, were performed after days of interactions (Gallardo-Navarro & Santillan, 2019; Piccardi *et al*., 2019; Mickalide & Kuehn, 2019). Considering that the generation time of most aerobic, free-living bacteria is 20 to 60 min under optimal conditions (i.e., those of experimental studies), the data from these experiments represent a late picture of an interaction that began within minutes of the bacteria meeting each other. In this context, there is a need for models that allow us to study the immediate response to competition in bacterial communities.

In previous work, after studying a transitive interaction network of 78 bacterial strains, a repeated pattern of Resistant (R), Antagonist (A), and Sensitive (S) strains was observed that reflected a history of interactions in communities. (Pérez-Gutiérrez *et al*., 2013). We constructed and evaluated a Bacillota (formerly Firmicutes) three-species synthetic community from this collection. We describe this synthetic community, which we named BARS (Bacillota Antagonist + Resistant + Sensitive). Each strain has a distinct colony phenotype that allows simple quantification of the viability of each strain in the interactions. The BARS community allowed us to observe emergent properties within minutes of the interaction, making it a unique model for exploring the immediate response to antagonism. BARS properties reproduce those of ecological communities: The effect that a strong competitor has on the activity of its interactor, habitat modification, and changes in the abundance of the sensitive strain. BARS represents a dynamic model for the study of bacterial interactions in a synthetic community with a simple experimental approach but still with properties of a complex system that permits the exploration of relevant properties and dynamics.

## 2 Materials and Methods

**Strains and media.** *S. horikoshii* CH20a (*Sh*20a), *B. pumilus* CH145 (*Bp*145) and *B. cereus* CH111 (*Bc*111) were reported previously by Pérez-Gutiérrez *et al*. (2013) and were isolated from the Churince system in the basin of Cuatrociénegas, Coahuila, México. The strains were propagated in Marine Medium agar (MMs) and growth in liquid Marine Medium (MMl) at 28° C with constant shaking. Marine Medium composition can be found in Cerritos *et al*. (2011).

### Generation of the interaction dynamics of the BARS community

Three strains were selected that belonged to different species in the Bacillota phylum that had distinct phenotypes on Marine Medium agar plates. To determine the ideal proportion of each strain for the interaction assays, the cell density of each strain was evaluated by confrontations between the strains directly in MMs. To perform the interaction assays, overnight cultures were used (14 h). For the start of growth, 100 μl of the overnight culture was transferred to 30 ml fresh medium in 250 ml nephelometric flasks, and growth was monitored using absorbance on the Klett Somerson colorimeter (red filter). Growth was monitored up to the exponential phase, and cultures were used in this phase. The interaction assays were started with a volume of 20 ml of *S. horikoshii* CH20a with approximately 1.5×10^9^ CFU/ml, 4 ml of *B. pumilus* CH145 with approximately 8×10^8^ CFU/ml and 4 ml of *B. cereus* CH111 with approximately 2×10^8^ CFU/ml. The interacting strains were cultured in 50 ml Falcon tubes, in a final volume of 40 ml. MMl was used to complete the volume of 40 ml for each interaction. Monocultures and different paired combinations and three strain combinations were monitored. Samples were obtained every 5 minutes beginning at time 0 with a duration of the interaction typically up to 30 min. Each sample was diluted by serial dilution through 1 × 10^-6^ and the three highest dilutions were plated on MMs. The plated samples were incubated at 28° C for 48 h. CFU counts for each colony phenotype were obtained (*Sh*20a yellow, *Bp*145 white, *Bc*111 large white colonies). All interactions were tested using data from at least three biological replicates. Changes in cell density or volume are explained for each experiment.

### Interaction experiments through membranes

For the interaction assays without cellular contact we followed the same methodology described above for the growth kinetics. The difference in these assays was the use of dialysis membranes to evaluate interactions only through diffusible metabolites (Master thesis Ortega-Aguilar, 2020). The membranes used were Spectra/Pro^®^ of 20 kDa pore. One ml of the *Sh*20a cell culture +/− 200 ul of either *Sh*20a or *Bp*145 was placed in a membrane and this was submerged in a 250 ml beaker with another cell culture (150 ml) and incubated at room temperature (25° C). The interaction was allowed to continue for 60 min to enable diffusion through the membrane. Samples were taken every 10 min, and used to prepare dilutions through 1 × 10^-6^ and the three highest dilutions were plated on MMs agar. The plated samples were incubated at 28° C for 48 h and the CFUs of each of the three colonial phenotypes were counted. At least three biological replicates were done for each interaction.

### Obtention of supernatants and lysates of *B. cereus* CH111

To obtain culture supernatants we followed the same procedure for culture conditions as in the interaction assays, at the same density of cells (2×10^8^ CFU/ml) and at the same point in the growth curve. Supernatants were centrifuged with a Sorvall Legend Micro 21R for 10 min at 13 rpm at room temperature, and separated and filtered with a 45 μm pore Millex□-GV membrane of PVDF (polyvinylidene difluoride) and nitrocellulose. For the experiments we substituted the culture of *B. cereus* CH111 with its supernatant and performed the experiment for the same 30 min with the strains *S. horikoshii* CH20a and *B. pumilus* CH145. Samples were obtained every 5 min, and dilutions and plating were done as described above.

To obtain a CH111 lysate, we used the same conditions for the other cultures of *B. cereus* CH111 and heated the culture for 30 min at 85 °C to lyse the cells. The lysate was used as part of an assay with the strains *S. horikoshii* CH20a and *B. pumilus* CH145 for 30 min and samples were taken every 5 min. Dilutions and plating were done as described above.

## 3 Results

### Design of the BARS synthetic community

In a previous study 78 sediment strains from a lagoon were tested in paired interactions for antagonism. These strains are mainly members of the phylum Bacillota, and their interaction network exhibited a transitive pattern where strains were either antagonistic, resistant, or sensitive (Perez *et al*. 2013). Their evaluated interactions within and across the communities from which they were isolated suggested a history of interaction. We built a small community by combining three strains that each exhibited a different trait. The BARS synthetic community comprises the resistant strain *Bacillus cereus* strain CH111 (*Bc*111), the antagonist strain *Bacillus pumilus* CH145 (*Bp*145), and the sensitive strain *Sutcliffiella horikoshii* CH20a (*Sh*20a) (it has recently been proposed that strains from *Bacillus horikoshii* be renamed *Sutcliffiella horikoshii* (Gupta *et al*. 2020)). As for the expected roles of the selected strains, resistant strain *Bc*111 exhibited no antagonistic capacity and was resistant to antagonism by 74 of the 78 strains evaluated (out-degree 0/in-degree 4); the antagonist strain *Bp*145a (out-degree 36/in-degree 4) was able to antagonize half of the tested strains; and the sensitive strain *Sh*20a was antagonized by 18 different strains (out-degree 0/in-degree 18). The three strains can grow in mono-culture in Marine Medium with doubling times of 38 min (*Sh*20a), 51 min (*Bp*145) and 30 min (*Bc*111). They have different colony phenotypes that allowed us to differentiate and count them in co-culture on semisolid agar plates (Supplementary Fig. 2).

### Higher-order interactions in the antagonist BARS dynamics

We used liquid cultures to evaluate the interaction among strains in a well-mixed environment. The main goal of our study was to monitor the dynamics of the interactions, but especially to evaluate the immediate response, that is, during 30 min, before the bacteria would begin cell division. Cultures from strains of *Bp*145 and *Sh*20a were grown as monocultures to exponential phase and then combined. Samples were taken at different times and plated. The proportion of *Sh*20a to *Bp145* at the start of the interaction was approximately 10:1 (1.5 × 10^9^ CFU/ml and 8 × 10^8^ CFU/ml, respectively). Figure 1 shows the CFU of each strain at different sampling times. The viable cell counts of *Sh*20a were reduced by more than 60% in the first 5 min of interaction with *Bp*145 (counts decreased to 5×10^8^ CFU/ml) (Fig. 1A). The reduction continued, but at a markedly lower pace, throughout the next 15 minutes, and no more decreases in CFUs were observed after 20 min. By 30 min, approximately 85% of the starting cell count of *Sh*20a was not viable (total reduction to 2×10^8^ CFU/ml). When paired interactions were set up in the same fashion, between *Sh*20a and *Bc*111 and between *Bp*145 and *Bc*111, no antagonism was observed and all strains maintained the same initial viability after 30 min (Figure 1C).

**Figure 1.**
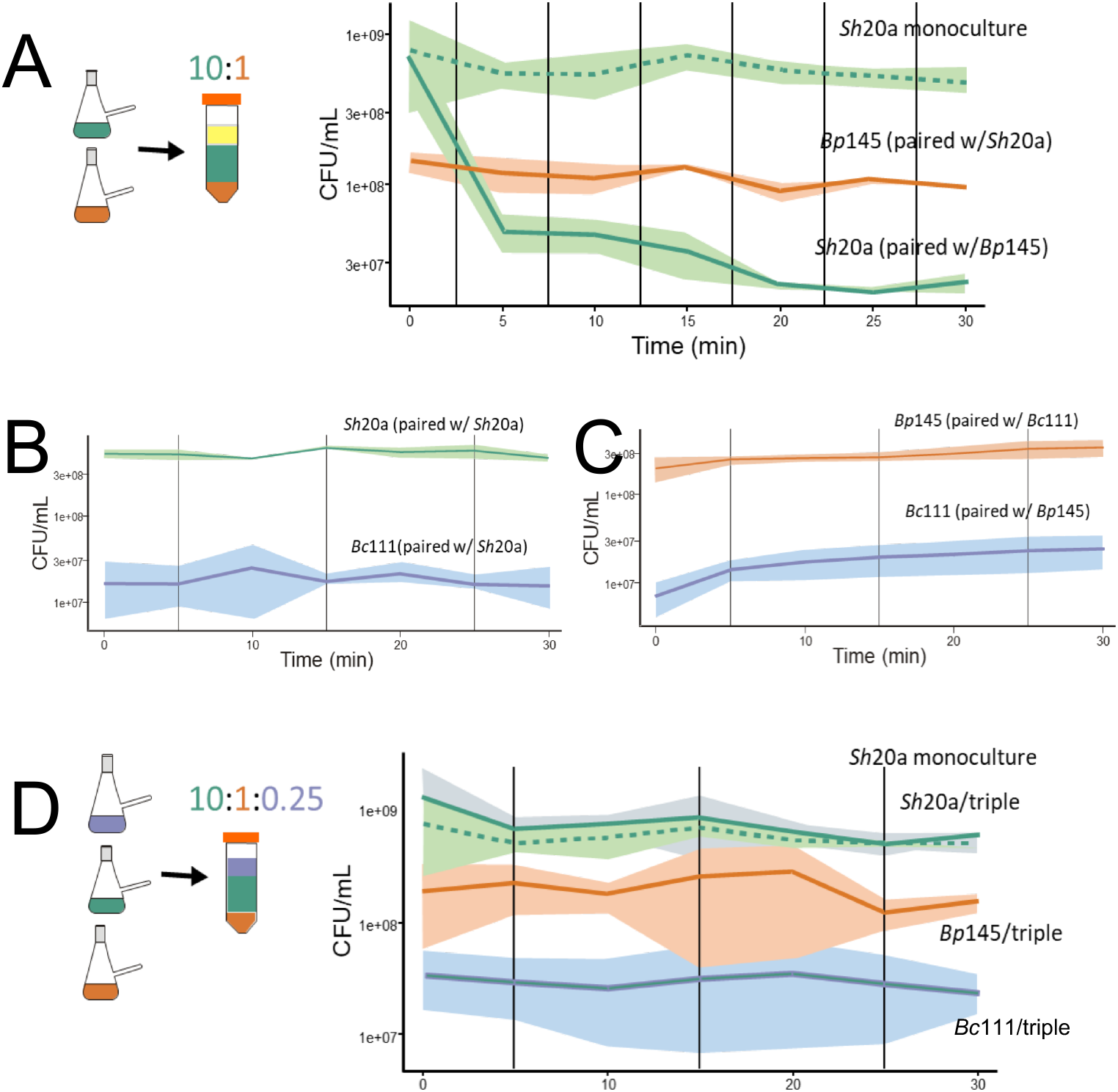
BARS synthetic community, from antagonistic paired interaction to emergent properties in a triple interaction. Viable cell counts over 30 minutes as measured by colony forming units (CFU). The viable count of each strain was obtained by plating at different time points after inoculating or combining monocultures; colonies of each phenotype were counted. Shadowed areas show the dispersion of data from three replicate experiments. **A.** Kinetics of combined sensitive strain *Sh*20a (green solid line) and antagonist strain *Bp*145 (orange solid line) shows an immediate antagonistic effect of strain *Bp*145 over strain *Sh*20a. The proportion of *Sh*20a:*Bp*145 was 10:1 in the paired interaction. *Sh*20a maintained in monoculture (green discontinuous line) is shown as a control. **B.** Kinetics of a paired interaction of resistant strain *Bc*111 (blue solid line) and sensitive strain *Sh*20a (green solid line). **C.** Kinetics of the combined antagonist strain *Bp*145 (orange solid line) and resistant strain *Bc*111 (blue solid line). **D.** No antagonism occurred in the triple interaction: resistant strain *Bc*111 (blue solid line), antagonist strain *Bp*145 (orange solid line), and sensitive strain *Sh*20a (green solid line). CFUs of *Sh*20a in monoculture are shown (green dotted line). In the triple interaction the ratio *Bh*20a:*Bp*145:*Bc*111 was 10:1:0.25. Tubes represent the combinations of strains in the interactions. Yellow represents culture medium; green, orange and blue represent monocultures from sensitive, antagonist and resistant strains.

The three strains were next combined and their viabilities were evaluated by plating at different times. *Sh*20a cell counts were not reduced despite the presence of the antagonist strain *Bp*145 (Fig 1C). The unexpected survival of *Sh*20a in interaction with antagonist *Bp145* is an emergent property that resulted from the presence of *Bc*111 in the interaction. The resistant strain seemed to neutralize the antagonistic interaction. This “stability” could be observed during the 30 minutes of the interaction. All strains maintained their CFUs counts when in monoculture (data not shown, except for *Sh*20a in Fig 1A).

### Antagonism without cellular contact

Microbes can harm each other directly through physical contact (i.e., toxin injection) or by secreting metabolites in the environment. To determine if cellular contact was needed for *Bp*145 to antagonize *Sh*20a, or if antagonism occurred through a diffusible molecule(s), we set up an experiment with dialysis membranes that permitted the exchange of metabolites and prevented cell-cell contact. The strains were grown to exponential phase and *Sh*20a was placed into membrane tubing (Spectra/Pro^®^ 20 kDa) that was introduced into a flask containing the antagonist strain *Bp*145. Antagonism between *Bp*145 and *Sh*20a was observed despite membrane separation (Supplementary Figure 3). The inhibition of the sensitive strain was observed from the beginning, as in the first 10 minutes the population of *Sh*20a decreased from 5.4 × 10^9^ CFU/ml to 1.1×10^9^ CFU/ml. No further inhibition occurred in the next 50 min (the duration of the assay was extended to account for a possible diffusion delay). Thus, antagonism *Bp*145 to *Sh*20a seemed to occur through diffusible molecule(s) and did not require cell-cell contact.

### A change of state from sensitive to tolerant occurred after 30 min as a result of the antagonistic interaction

We wondered why the sensitive strain did not suffer further loss of viability after 20 min in paired interaction with the antagonist strain. Fresh *Sh*20a strain culture, in exponential phase, was added 30 min after a *Sh*20a/*Bp*145 interaction had been initiated, and the interaction was allowed to take place for 30 minutes. As observed in Figure 2, in this second phase of interaction, the viability (counted as CFU) of strain *Sh*20a did not decrease. Bp145 did not antagonize the freshly added Sh20a.

**Figure 2.**
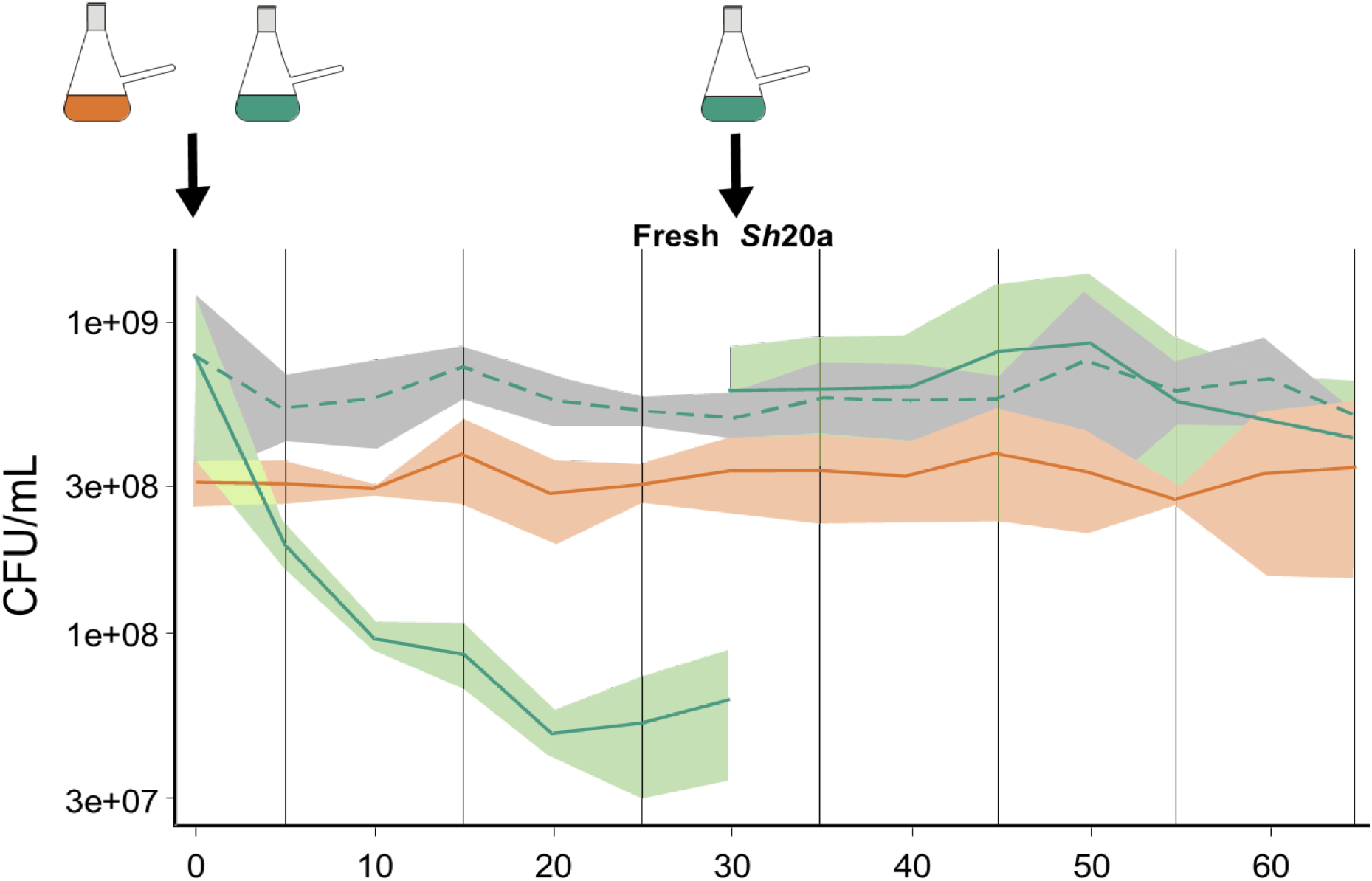
Change of state in the antagonism dynamics between sensitive and antagonist strains. The first 30 min show the CFU counts of each strain in the paired interaction between *Sh*20a strain (green solid line) with *Bp*145 strain (orange solid line) where the expected decrease in CFU due to antagonism was observed. At 30 min, fresh *Sh*20a culture was added to the ongoing interaction and we monitored the viability of the strains for another 30 min. After this time, no change in the viable count was observed for the *Sh*20a strain (green solid line). As a control, the CFU of the *Sh*20a strain in monoculture (green discontinuous line) is shown.

Different hypotheses could explain this result: (1) detoxification of the media, such that an antagonist metabolite was neutralized or destroyed, (2) *Bp*145 changed to a non-antagonistic state, or (3) *Sh*20a immediately acquired tolerance after sensing dead cells of the old culture as a signal to induce tolerance. A combination of these situations could explain the result. We next carried out new experiments using the cells of *Sh*20a that survived the interaction with *Bp*145 and tested if pre-exposed cells of *Sh*20a could experience a change of state and modify their interaction with *Bp*145.

### Pre-exposure of *S. horikoshii* CH20a cells to the antagonist provides resistance to a second challenge

That *Sh*20a survived the antagonism with strain *Bp*145 could be explained by a population of “persister cells”, defined as cells in a susceptible population that tolerate antagonistic substances (i.e., antibiotics) (Levin & Rozen, 2004). Another possibility is that contact with the antagonist could have induced tolerance to the antagonistic molecule. We evaluated whether *Sh*20a could change its state from sensitive to resistant after exposure to the antagonist *Bp*145 strain. We set up an experiment with dialysis membranes, which allowed metabolic exchange while preventing contact between sensitive and resistant cells. Using cells grown in monoculture to exponential phase, *Sh*20a was placed inside the membrane and *Bp*145 outside the membrane. The first part of the interaction took place for 60 minutes (up from 30 min compared to previous experiments to allow time for the diffusion of metabolites across the membrane), and the dialysis bags containing *Sh*20a cells were then transferred to a flask containing a new culture of *Bp*145, for another 60 minutes. Figure 3A shows that the *Sh*20a cells that had been exposed to *Bp*145 did not show the dramatic loss of viability that was typically exhibited by unexposed cells upon incubation with a fresh culture of *Bp*145. This result supports our hypothesis that, in *Sh*20a, tolerance to antagonism was induced by exposure to *Bp*145.

**Figure 3.**
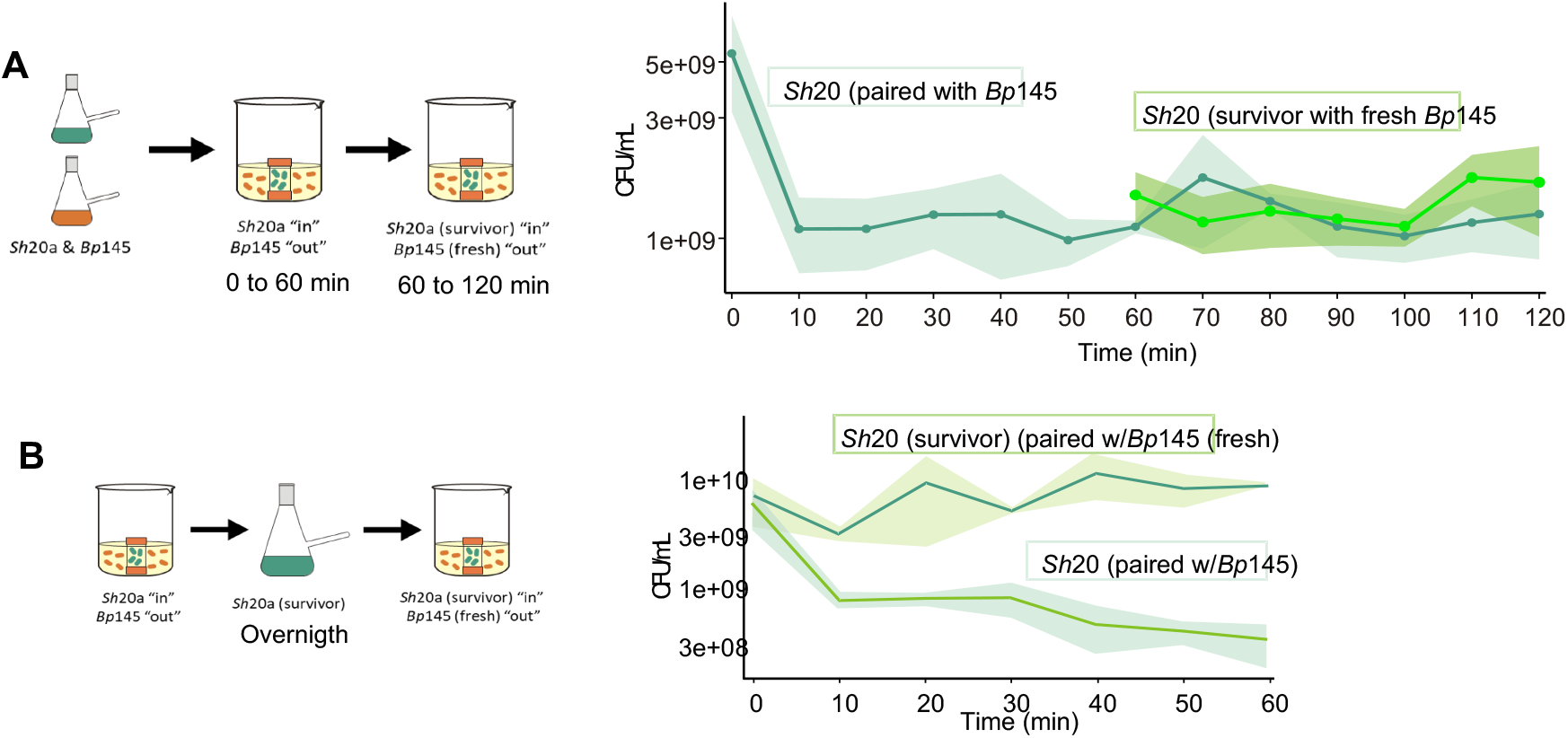
Acquired tolerance of the *Sh20a* sensitive strain to antagonism from *Bp145*. **A.** Dynamics of sensitive cells that survived antagonism in a second interaction with *Bp*145. A culture of *Sh*20a was introduced into membranes that were exposed to the antagonist strain *Bp*145. It can be observed that the CFUs of *Sh*20a in this first interaction with *Bp*145 (green solid line) decreased in the first 10 minutes, but their viability did not change further through the next 50 min. After 60 min, the membranes containing the *Sh*20a (surviving cells) were transferred to another beaker that contained fresh cells of antagonist *Bp*145. The counts of these survivor cells of *Sh*20a (light green line) did not decrease further by interaction with fresh *Bp*145. The control cells in membranes that were not transferred maintained the CFU counts (green solid line). **B.** Dynamics of *Sh*20a cells that survived antagonism and were grown in monoculture for 14 h overnight and then subjected to a second interaction with the antagonist strain. In the first interaction, the CFUs of the *Sh*20a cells decreased (continuous green line). In the second interaction, the counts of *Sh*20a survivor cells did not decrease upon exposure to *Bp*145, and thus *Sh*20a survivor cells tolerated a new confrontation with the antagonist.

To explore if survivor cells of *Sh*20a that became tolerant to *Bp*145 could maintain tolerance after cell division, survivor cells of *Sh*20a were grown overnight and used as inoculum to start a culture that was grown to exponential phase. These growing cells were placed in a membrane and submerged in a fresh culture of *Bp*145 (Figure 3B). The pre-exposed-and-regrown cells of *Sh*20a showed higher tolerance than unexposed cells. The antagonism of *Bp*145 was reduced compared with an interaction between a fresh culture of *Sh*20a and *Bp*145 (Figure 3B). One explanation for the partial survival of *Sh*20a exposed to *Bp*145 could be that the concentration of the antagonistic compound was not high enough to kill the entire population, which allowed part of the population to express genes that conferred this tolerance and that this tolerance phenotype was maintained even after overnight growth. The presence of persister cells from the beginning of the experiment could thus be ruled out because if the pattern were due to persister cells, we would have expected 60% of cells to have been sensitive after a regrowth, whereas in this new experiment, only 20% of cells were sensitive (Figure 3B) (from 1.4 × 10^10^ CFU/ml to 1 × 10^10^ CFU/ml).

### Live cells of *Bc111* are more effective in the neutralization of antagonism than its supernatant and cell lysates

We evaluated whether a lysate of *Bc*111 (prepared by heat treatment of cells at 85 °C) could protect sensitive *Sh*20a cells from antagonism. The lysate of *Bc*111 did not eliminate the antagonistic effect of *Bp*145 over *Sh*20a, as 75 % of *Sh*20a cells were antagonized (from 1.5×10^9^ CFU/ml to 3.5×10^8^ CFU/ml) (Supplementary Fig. 4). It is possible that a metabolite produced by *Bc*111 was heat denatured during the treatment to obtain the culture lysate and therefore the lysate could not fully protect. Thus, we also evaluated whether filtered (0. 45 μm Millex^®^-GV filter) *Bc*111 supernatant could neutralize antagonism. Adding the supernatant showed that the population of *Sh*20a cells decreased at a slower rate compared to the interaction between only *Sh*20a and *Bp*145 (from 1.5 x10^9^ CFU/ml to 2.1×10^8^ CFU/ml) (Supplementary Fig. 4). These results suggested that although the supernatant from *Bc*111 somewhat neutralized the antagonistic substance, only the live *Bc*111 strain conferred the full protection observed in the triple interaction. These results indicate that the mechanism of protection of the resistant strain is a combination between molecules already expressed and found in the lysate, and the expression of a possible metabolite during the interaction.

### The dynamics of the triple interaction are highly sensitive to the abundance of the resistant strain

We evaluated the importance of the proportions of sensitive, antagonistic, and resistant strains in the interaction. The proportion of cells in the BARS interactions used until now had been 10:1:0.25 (sensitive: antagonist: resistant, respectively). We next tested different proportions changing only the abundance of the resistant strain (proportions tested: 10:1:0.15; 10:1:0.075; 10:1:0.05). Reducing the proportion of *Bc*111 to 0.15 (1.3×10^8^ CFU / ml) still protected the sensitive strain but when the concentration of *Bc*111 was lowered to 0.075 (6.6×10^7^ CFU/ml), protection decreased (Figure 4). However, when the cells of *Bc*111 were further decreased to 10:1:0.05 (4×10^7^ CFU/ml), an unexpected negative effect was observed on the population of *Sh*20a. The sensitive strain was antagonized even more drastically than in the antagonism observed in the paired interaction between *Sh*20a and *Bp*145. These results suggest that the cell density of *Bc*111 determines the fate of *Sh*20a when exposed to the antagonism of *Bp*145.

**Figure 4.**
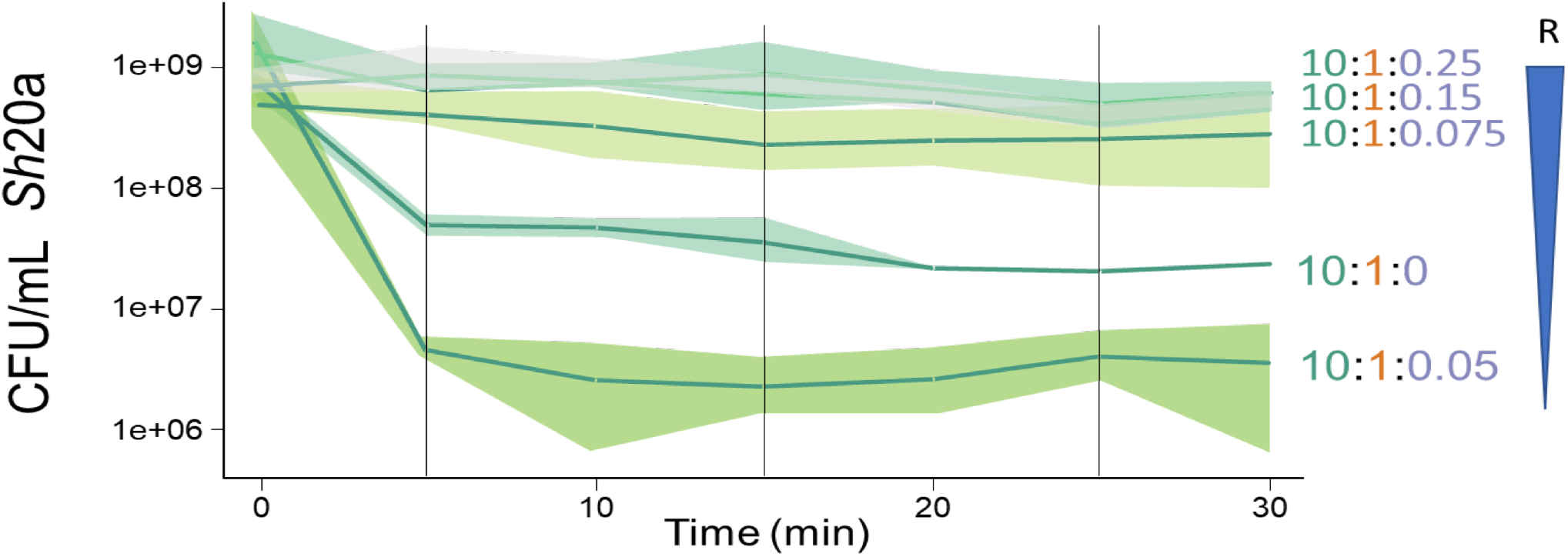
The dynamics of the Sh20a strain is highly sensitive to a decrease in the proportion of Bc111 in a triple interaction. A reduction in the proportion of Bc111 relative to the sensitive and antagonist cells in triple interaction (schematized by the inverted blue triangle) affects the survival of Sh20a. At 2×10^8^ CFU/ml the sensitive strain is not antagonized in the presence of Bp145 (dark blue line). Upon sequential reductions from 1.3×10^8^ CFU/ml (10:1:0.6) and 6.6×10^7^ CFU/ml 10:1:0.3), a slight decrease of the population of the Sh20a is observed. Unexpectedly, a reduction to 4×10^7^ CFU/ml results in a large reduction in the population of Sh20a (10:1:0.02), an effect even more dramatic than in a paired interaction (10:1:0).

### The BARS model and its properties as a higher-order system

The strains in the BARS model (Fig. 5) derive from a collection of 78 strains, which, when evaluated in pairs for antagonism, each fell in a hierarchical network that allowed them to be classified either as resistant, antagonist, or sensitive. Traits were commonly shared between taxonomically similar strains (Perez et al. 2013). In Perez et al. (2013), none of the traits could be deduced by studying each strain in isolation, and in setting up the BARS model, we show that their pairwise interaction also did not allow for predictions of their behavior as a triad. Another feature of the BARS model is the response to interaction, which can be evaluated within 30 min, before cell division takes place. This is relevant because in such a short time the culture medium has not been depleted so the antagonistic response can be isolated from resource competition. Our model shows that part of the population of the sensitive strain survived antagonism and also seemed to be able to induce tolerance in minutes. Also, within minutes, the *Bc*111 strain promoted survival of the sensitive strain even in the presence of the antagonistic strain.

**Figure 5.**
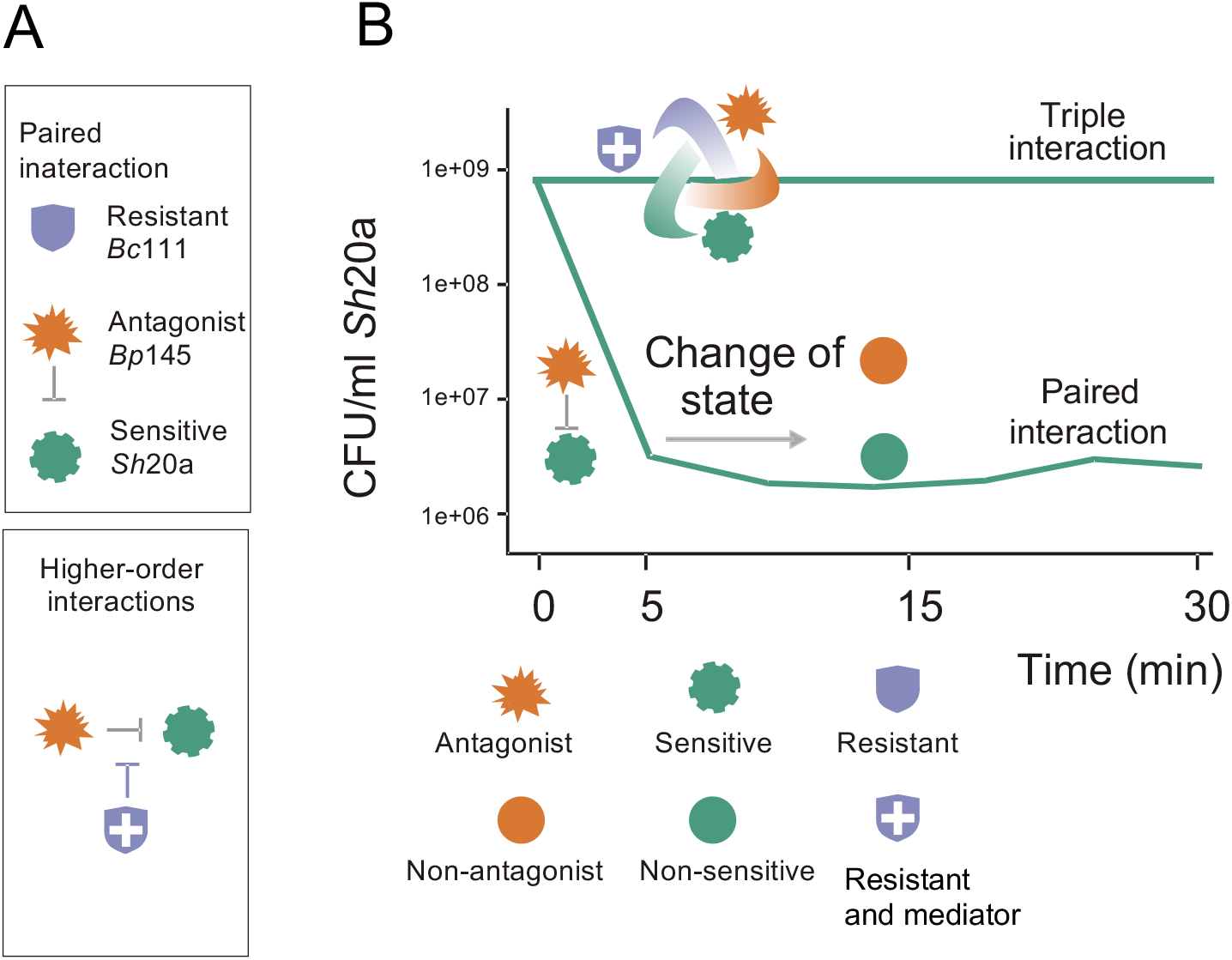
Immediate response dynamics and emergent properties in the three-strain BARS synthetic community model. A. Each strain in the BARS model exhibits known properties from paired interactions that do not, however, enable the prediction of their behavior in a triple community. As a higher-order interaction system, it adopts features of complex communities where *Bc1*11 seems to act as a mediator (blue shield symbol). B. In a paired interaction, a strong antagonism is observed in the first five minutes, but then no further antagonism is observed in the next 20 min (shown by the dark green curve that decreases and then remains stable). A change of state of strains *B*p145, represented by symbols changing from spiked to round, is inferred from experiments showing that after 30 min interaction with strain Sh20a, it does not antagonize “fresh” strain *Sh*20a. Similarly, *Sh*20a that survived antagonism from *Bp*145 becomes resistant to antagonism when exposed to “fresh” *Bp*145. Change from sensitive to non-sensitive (green wavy symbol to round symbol). The top line (light green) shows that a stable three-strain community arises as an emergent property, and antagonism is not observed. As an emergent interaction, we do not know what type of relations are established between strains and/or the environment. We show that antagonism is mediated by diffusible metabolites and is cell contact independent. The triple interaction dynamics is sensitive to changes in the proportion of the resistant strain.

One of the most critical features of the BARS model is that the triad achieved a stable coordinated system immediately after being placed in interaction after strains had been grown as monocultures. This state could not be predicted from the paired interactions, and was an emergent state, which is a property of higher-order interactions.

## 4 Discussion

In this work, we present a synthetic community that we denominated BARS, constituted of three Bacillota strains that have different characteristics: antagonist (A), resistant to antagonism (R), and sensitive to antagonism (S). The criteria for selecting these strains was that they had different ecological roles and colony phenotypes. The roles of these three strains were known from our previous work, where paired interactions in a 78 × 78 strain matrix were evaluated (Peréz-Gutiérrez *et al*., 2013). In the resulting network, these three strains clearly differed in their antagonism and formed non-hierarchical interactions. It is interesting to contemplate that what we selected for in the triad were not random strains but strains with a history of community interactions that evolved adaptive responses to complex interactions. The genetic repertoire for exerting antagonism, inducing resistance, and the capacity to detoxify, have all been acquired and fine-tuned through their evolution in communities.

The importance of BARS and other models in synthetic ecology is that they allow us to address questions regarding the assembly of communities, a process where biotic and abiotic filters define the diversity and abundance of strains. Given the complexity of this endeavor, having models that reduce the number of individuals and environmental variables is essential. However, equally important and missing in most microbial models is being able to address the immediate consequence of the invasion of a community by new individuals. In the plot of microbial assembly, in most studies, we are probably missing the climax and seeing only the resolution, a point where some kind of negotiation between individuals has already occurred. BARS shows that within 30 min, that is, before the cells divide, the negotiation was influenced by different variables leading to changes of cell states.

Additionally, since we evaluated the antagonistic effect within the first minutes of the interaction, we probably are not observing resource competition; therefore, this model reduces yet another variable. Although physiological changes may occur upon transfer to fresh medium, since the same medium and temperature are maintained, and the cells are in log phase upon setting up the interactions, we suggest that the cells are not limited by nutrient availability and are not competing for resources.

The study of microbial interactions commonly considers the influence of spatial structure in cooperation dynamics. Spatial structure is suggested to be an essential factor for coexistence. Zapién-Campos *et al*. (2015) modeled the interaction of the 78 strains and reported the formation of patches where resistant strains protected sensitive strains from the antagonist strains and, in this way, determined the survival of individuals under antagonistic dynamics. We observed similar patches upon plating the BARS community (Supplementary Fig. 2). However, in a well-mixed environment, such as that of liquid cultures, the diffusion of molecules leads to unavoidable interaction between bacteria.

Many models evaluate interactions under homogeneous conditions to consider either cooperation or competition for resources (Reviewed in Hibbing *et al*., 2010). Spatial structure is thus another variable absent in the BARS model as our evaluations occurred in a well-mixed environment (liquid medium). Contrary to the classical rock-paper-scissors model, our model does not show a cyclic dynamic because *B. cereus* CH111 modified the interaction between *S. horikoshii* CH20a and *B. pumilus* CH145. However, in a triad model where the three strains coexisted, it was not possible to recapitulate the paired dynamics.

A few examples of HOI in synthetic communities of microorganisms have been reported in recent years (Gallardo & Santillan, 2019; Lozano *et al*., 2019; Piccardi *et al*., 2019; Mickalide & Kuehn, 2019), some of them showing some properties similar to the BARS community dynamics. Gallardo-Navarro & Santillan (2019) described a model in which a resistant strain (*Staphylococcus sp.*) seems to protect a sensitive one (*B. aquimaris*) from the antagonism of a *B. pumilus* strain. Another example comes from the work of Lozano *et al*. (2019), in a model named THOR (the hitchhikers of the rhizosphere), with a similar three-species community; in this model, the resistant strain is also *B. cereus* that protects a strain of *Flavobacterium johnsoniae* from growth inhibition by *Pseudomonas koreensis*. Given the emergent properties that we and others have observed, maybe we have to carefully evaluate more data before applying a “rule” such as the one proposed by Friedman for the prediction that species that all coexist with each other in pairs will survive, whereas species that are excluded by any of the surviving species will go extinct (Friedman *et al*., 2017).

Since *Bc*111 seems to mediate the ability of the three strains to coexist, and, apparently, even its supernatant offers some protection, we speculate that it can neutralize or detoxify the environment from the antagonist substance produced by *Bp*145. The detoxification of the environment by one of the interactors in a community has been described. The idea of a third interactor as a mediator of antagonism was reported by Kelsic *et al*. (2015). They proposed that the antibiotic-resistant strains in a community have the function of degrading the toxins and antibiotics in the environment, allowing a sensitive strain to cohabit in the community with the antibiotic producer strains. In studies on antibiotic resistance, detoxifier capacities have been attributed to strains able to express enzymes that can degrade antibiotics, such as β-lactamases. In a gram-negative model the presence of β-lactamases was reported as a modulator of an antagonistic interaction by the cleavage and deactivation of antibiotics (Medaney *et al*., 2016; Frost *et al*., 2018). In BARS, the ratio *Bc*111 relative to the antagonist and sensitive strains defines whether the antagonistic interaction is subdued or increased. In the report from Medaney *et al*. (2016), they observed an increase in the cell density of the interactors as an emergent property, since the population of the sensitive strain was positively correlated with the cell density of the resistant one. An important difference with the BARS model, is that at a lower cell density of the resistant strain, the sensitive *Sh*20a seemed to be more affected by the antagonist. Abrudan *et al*. (2015) used a *Streptomyces* system to evaluate paired growth and observed that in some cases, a third interactor could increase the antagonism between two interactors.

Many strategies for antagonism have been reported, such as the secretion of enzymes or metabolites directly to the extracellular space or even through outer membrane vesicles (OMVs). The production of OMVs protects against harmful substances such as reactive oxygen species, antibiotics, antimicrobial peptides and phages (Lekmeechai *et al*., 2018; Liao *et al*., 2015; Kulkarni *et al*., 2015; Manning and Kuehn, 2011; Roszkowiak *et al*., 2019). It is important, however, to consider that antagonistic molecules also have other functions, and not just antagonism. It has been described that antibiotics are likely to be found in the environment at low concentrations, and that they may play important roles in the modulation of metabolic function in natural microbial communities (Yim *et al*., 2006). Although toxin production is a common response for competitive interactions, there is a wide range of responses and it has been observed that factors such as strain frequency, nutrient level, the level of strain mixing (relatedness), and the cost of toxin production are all important for the bacterial decision to produce toxins for an interaction (Niehus *et al*., 2021).

Tolerance of a fraction of the *Sh*20a population to antagonism by *Bp*145 is a property of the dynamics of the BARS community. *Sh*20a seems to become tolerant during its interaction with *Bp*145 but there is the possibility that a subpopulation of *Sh*20a was already tolerant.

Communities experience invasion continuously, or their members get shuffled around to find themselves with new neighbors. They have evolved adaptations to respond immediately to biotic and abiotic challenges in complex communities, making them experts in integrating fine-grained information and collectively finding solutions that we observe as emergent phenomena. Their capacity to respond immediately was probably shaped by the history of past environmental states. As Flak states, “Biological systems can be viewed as a hypothesis about the present and future environments they or their offspring will encounter, induced from the history of past environmental states they or their ancestors have experienced” J. Flack (2017). We expect that the BARS model is simple enough to allow deep exploration of the molecular mechanisms underlying the changes of state and a fast response when in paired interaction. Through genomic and transcriptomics analysis, it may be possible to identify genes involved in the immediate response to community interactions as well as the specific environmental and biotic cues that trigger such a response. This information will aid in modeling the changes of state in the dynamics of the BARS community. The principles we can gather from studying such a model will hopefully provide clues on how other communities respond to antagonistic interactions.

## 5. CONCLUSION

The BARS three-strain synthetic community offers a good HOI model that exhibits emergent properties. As communities are complex systems, we cannot predict what will happen when more than two individuals interact. At higher-order levels, emergent properties arise, as in the case of BARS, where antagonistic competition is no longer observed. Thus, the BARS model could be excellent for studying natural bacterial communities and a good example of HOI in bacterial ecology.

Of great significance is that the BARS model allows for the evaluation of the immediate response of bacteria in antagonistic competition. Most studies evaluate community assembly in hours or days. In these settings, we do not know what changes were experienced by its members upon a first encounter. This approach makes the community assembly process a black box; we miss the game’s climax and are left only with a picture of the winners. The BARS synthetic community simplifies the natural interactions but maintains important features of ecological communities.

## 6. Acknowledgements

This project was financed by Conacyt Ciencia de Frontera No. 39587 to G.O.A. B.A.S. acknowledges fellowship from Conacyt. We acknowledge MC Africa Islas-Robles, Maricruz Gonzalez Alvarado, members of the Olmedo-Alvarez group for helpful discussions. We thank José Antonio Cisneros for his help with photographs and Jorge Eugenio Reynoso for his help with the drawings. We thank Mike Travisano and Martin Heil for fruitful discusions.

---

**Supplementary Figure 1.** Strain selection for BARS. Based on a past evaluation by Pérez-Gutiérrez *et al*. (2013), paired interaction phenotypes were re-evaluated on a mat from a *Sutcliffiella horikoshii* 20a culture. Strain *Bacillus pumillus* 145 was selected as the antagonist strain and *Bacillus cereus* 111 as a non-antagonist and resistant strain (resistance was evaluated in a separate experiment).

**Supplementary Figure 2.** Phenotypes of BARS strains. A. All strains were grown on Marine Medium. Shown are their duplication times, all below 60 min. B. The out-degree and in-degree phenotype data were obtained from the paired interaction experiments of the 78 × 78 matrix described in Pérez-Gutiérrez *et al* (2013). In-degree is the number of strains that antagonized a given strain, and out-degree is the number of strains that a given strain antagonized. The BARS strains exhibited different colony morphologies that allow their quantification upon plating dilutions of combined cultures. Plates in C show colonies after 5 min of paired interaction between antagonist strains *Bp*145 and *Sh*20a. Zooming in allows us to more easily see the different phenotypes of the colonies, and particularly the survival of sensitive (yellow) only in patches away from the antagonist (whitish). Plates in D show the results after 5 min of interaction between antagonist strains *Bp*145 (white, medium-sized colonies), *Sh*20a (yellow), and Bc111 (large white colonies).

**Supplementary figure 3. Dynamics of the BARS interaction can be reproduced through a membrane and are thus independent of cell-cell contact. A.** Antagonism in the BARS model occurs when cells of different strains are separated by a membrane, possibly through diffusion of metabolites across the membrane. Sensitive strain *Sh*20a was placed inside the membrane and placed in a beaker containing the *Bp*145 culture. Aliquots from each strain were plated at different times to determine CFU counts. The population of *Sh*20a (green solid line) was antagonized by *Bp*145 (red solid line) without the need for cell contact. **B.** *Bc*111 was able to stabilize the antagonistic interaction without cell contact. *B. cereus* 111, was placed out of the membrane (blue solid line), and *Sh*20a (green solid line) and *Bp*145 (red solid line) were both inside the membrane.

**Supplementary figure 4. Effect of *Bc*111’s lysate or supernatant on an antagonistic interaction. A.** Addition of a lysed culture of the resistant *Bc*111strain to a paired interaction between *Bp*145 (orange solid line) and *Sh*20a (green solid line), reduced antagonism. *Sh*20a in monoculture (no lysate added) was evaluated as a control (green discontinuous line). **B.** A filtered supernatant from a *Bc*111 culture was added to an interaction experiment where the antagonist *Bp*145 (orange solid line) and sensitive *Sh*20a (green solid line) were combined. The supernatant reduced and retarded the antagonism of *Bp*145 over *Sh*20a. *Sh*20a in monoculture (and no supernatant added) is shown as a control (green discontinuous line).

## References

Abrudan, M. I., Smakman, F., Grimbergen, A. J., Westhoff, S., Miller, E. L., van Wezel, G. P., & Rozen, D. E. (2015). Socially mediated induction and suppression of antibiosis during bacterial coexistence. Proceedings of the National Academy of Sciences, 112(35), 11054–11059. https://doi.org/10.1073/pnas.1504076112

Billick, I., & Case, T. J. (1994). Higher Order Interactions in Ecological Communities: What Are They and How Can They be Detected? Ecology, 75(6), 1529–1543. https://doi.org/10.2307/1939614

Carrara, F., Giometto, A., Seymour, M., Rinaldo, A., & Altermatt, F. (2015). Inferring species interactions in ecological communities: a comparison of methods at different levels of complexity. Methods in Ecology and Evolution, 6(8), 895–906. https://doi.org/10.1111/2041-210X.12363

Cerritos, R., Eguiarte, L. E., Avitia, M., Siefert, J., Travisano, M., Rodríguez-Verdugo, A., & Souza, V. (2011). Diversity of culturable thermo-resistant aquatic bacteria along an environmental gradient in Cuatro Ciénegas, Coahuila, México. Antonie van Leeuwenhoek, International Journal of General and Molecular Microbiology, 99(2), 303–318. https://doi.org/10.1007/s10482-010-9490-9

Cornforth, D. M., & Foster, K. R. (2013). Competition sensing: The social side of bacterial stress responses. Nature Reviews Microbiology, 11(4), 285–293. https://doi.org/10.1038/nrmicro2977

Flack, J. (2017). Life’s Information Hierarchy. In S. Walker, P. Davies, & G. Ellis (Eds.), From Matter to Life: Information and Causality (pp. 283–302). Cambridge: Cambridge University Press. doi:10.1017/9781316584200.012

Friedman, J., Higgins, L. M., & Gore, J. (2017). Community structure follows simple assembly rules in microbial microcosms. Nature Ecology and Evolution, 1(5), 1–7. https://doi.org/10.1038/s41559-017-0109

Frost, I., Smith, W. P. J., Mitri, S., Alvaro, ·, Millan, S., Davit, Y., … Foster, K. R. (2018). Cooperation, competition and antibiotic resistance in bacterial colonies. The ISME Journal, 12, 1582–1593. https://doi.org/10.1038/s41396-018-0090-4

Gallardo-Navarro, Ó. A., & Santillán, M. (2016). Three-way interactions in an artificial community of bacterial strains directly isolated from the environment and their effect on the system population dynamics. Frontiers, 10(November), 1–13. https://doi.org/10.3389/fmicb.2019.02555

Garcia, E. C. (2018). Contact-dependent interbacterial toxins deliver a message. Current Opinion in Microbiology, 42, 40–46. https://doi.org/10.1016/J.MIB.2017.09.011

Grilli, J., Barabás, G., Michalska-Smith, M. J., & Allesina, S. (2017). Higher-order interactions stabilize dynamics in competitive network models. Nature, 548(7666), 210–213. https://doi.org/10.1038/nature23273

Gupta, R. S., Patel, S., Saini, N., & Chen, S. (2020). Robust demarcation of 17 distinct bacillus species clades, proposed as novel bacillaceae genera, by phylogenomics and comparative genomic analyses: Description of robertmurraya kyonggiensis sp. nov. and proposal for an emended genus bacillus limiting it only to the members of the subtilis and cereus clades of species. International Journal of Systematic and Evolutionary Microbiology, 70(11), 5753–5798. https://doi.org/10.1099/IJSEM.0.004475/CITE/REFWORKS

Hibbing, M. E., Fuqua, C., Parsek, M. R., & Peterson, S. B. (n.d.). Bacterial competition: surviving and thriving in the microbial jungle. https://doi.org/10.1038/nrmicro2259

Kelsic, E. D., Zhao, J., Vetsigian, K., & Kishony, R. (2015). Counteraction of antibiotic production and degradation stabilizes microbial communities. Nature, 521(7553), 516–519. https://doi.org/10.1038/nature14485

Kerr, B., Riley, M. A., Feldman, M. W., & Bohannan, B. J. M. (2002). Local dispersal promotes biodiversity in a real-life game of rock–paper–scissors. Nature, 418(6894), 171–174. https://doi.org/10.1038/nature00823

Kirkup, B. C., & Riley, M. A. (2004). Antibiotic-mediated antagonism leads to a bacterial game of rock-paper-scissors in vivo. Nature, 428(6981), 412–414. https://doi.org/10.1038/nature02429

Kulkarni, H. M., Nagaraj, R., & Jagannadham, M. V. (2015). Protective role of E. coli outer membrane vesicles against antibiotics. Microbiological Research, 181, 1–7. https://doi.org/10.1016/J.MICRES.2015.07.008

Lekmeechai, S., Su, Y. C., Brant, M., Alvarado-Kristensson, M., Vallström, A., Obi, I., … Riesbeck, K. (2018). Helicobacter pylori Outer Membrane Vesicles Protect the Pathogen From Reactive Oxygen Species of the Respiratory Burst. Frontiers in Microbiology, 9(SEP). https://doi.org/10.3389/FMICB.2018.01837

Levin, B. R., & Rozen, D. E. (2006). Non-inherited antibiotic resistance. Nature Reviews Microbiology 2006 4:7, 4(7), 556–562. https://doi.org/10.1038/nrmicro1445

Levine, J. M., Bascompte, J., Adler, P. B., & Allesina, S. (2017, May 31). Beyond pairwise mechanisms of species coexistence in complex communities. Nature, Vol. 546, pp. 56–64. https://doi.org/10.1038/nature22898

Liao, Y. T., Kuo, S. C., Chiang, M. H., Lee, Y. T., Sung, W. C., Chen, Y. H., … Fung, C. P. (2015). Acinetobacter baumannii extracellular OXA-58 is primarily and selectively released via outer membrane vesicles after sec-dependent periplasmic translocation. Antimicrobial Agents and Chemotherapy, 59(12), 7346–7354. https://doi.org/10.1128/AAC.01343-15/SUPPL_FILE/ZAC012154583SO1.PDF

Lyons, N. A., & Kolter, R. (2017). Bacillus subtilis protects public goods by extending kin discrimination to closely related species. MBio, 8(4), e00723–17. https://doi.org/10.1128/mBio.00723-17

Lozano, G. L., Bravo, J. I., Garavito Diago, M. F., Park, H. B., Hurley, A., Peterson, S. B., … Handelsman, J. (2019). Introducing THOR, a model microbiome for genetic dissection of community behavior. MBio, 10(2), 1–14. https://doi.org/10.1128/mBio.02846-18

Manning, A. J., & Kuehn, M. J. (2011). Contribution of bacterial outer membrane vesicles to innate bacterial defense. BMC Microbiology, 11. https://doi.org/10.1186/1471-2180-11-258

Medaney, F., Dimitriu, T., Ellis, R. J., & Raymond, B. (2016). Live to cheat another day: Bacterial dormancy facilitates the social exploitation of β-lactamases. ISME Journal, 10(3), 778–787. https://doi.org/10.1038/ismej.2015.154

Mickalide, H., & Kuehn, S. (2019). Higher-Order Interaction between Species Inhibits Bacterial Invasion of a Phototroph-Predator Microbial Community. Cell Systems, 9(6), 521–533.e10. https://doi.org/10.1016/j.cels.2019.11.004

Nemergut, D. R., Schmidt, S. K., Fukami, T., O’Neill, S. P., Bilinski, T. M., Stanish, L. F., … Ferrenberg, S. (2013). Patterns and Processes of Microbial Community Assembly. Microbiology and Molecular Biology Reviews, 77(3), 342–356. https://doi.org/10.1128/MMBR.00051-12/ASSET/64D2229D-8BE5-48FA-B20E-B8585AB15584/ASSETS/GRAPHIC/ZMR9990923330004.JPEG

Niehus, R., Oliveira, N. M., Li, A., Fletcher, A. G., & Foster, K. R. (2021). The evolution of strategy in bacterial warfare via the regulation of bacteriocins and antibiotics. ELife, 10. https://doi.org/10.7554/ELIFE.69756

Ortega-Aguilar, Jaime (2020) “Análisis de la dinámica de comunidades de Bacillus spp. con control intercambio de metabolitos y en mesocosmos”. Master thesis Cinvestav 2020.

Pérez-Gutiérrez, R. A., López-Ramírez, V., Islas, Á., Alcaraz, L. D., Hernández-González, I., Olivera, B. C. L., … Olmedo-Alvarez, G. (2013). Antagonism influences assembly of a Bacillus guild in a local community and is depicted as a food-chain network. ISME Journal, 7(3), 487–497. https://doi.org/10.1038/ismej.2012.119

Piccardi, P., Vessman, B., & Mitri, S. (2019). Toxicity drives facilitation between 4 bacterial species. Proceedings of the National Academy of Sciences of the United States of America, 116(32), 15979–15984. https://doi.org/10.1073/pnas.1906172116

Roszkowiak, J., Jajor, P., Guła, G., Gubernator, J., Żak, A., Drulis-Kawa, Z., & Augustyniak, D. (2019). Interspecies Outer Membrane Vesicles (OMVs) Modulate the Sensitivity of Pathogenic Bacteria and Pathogenic Yeasts to Cationic Peptides and Serum Complement. International Journal of Molecular Sciences, 20(22). https://doi.org/10.3390/IJMS20225577

Sanchez-Gorostiaga, A., Bajić, D., Osborne, M. L., Poyatos, J. F., & Sanchez, A. (2019). High-order interactions distort the functional landscape of microbial consortia? In PLoS Biology (Vol. 17). https://doi.org/10.1371/journal.pbio.3000550

Sharpe, M. E., Hauser, P. M., Sharpe, R. G., & Errington, J. (1998). Bacillus subtilisCell Cycle as Studied by Fluorescence Microscopy:Constancy of Cell Length at Initiation of DNA Replicationand Evidence for Active Nucleoid Partitioning. In JOURNAL OF BACTERIOLOGY (Vol. 180). Retrieved from https://journals.asm.org/journal/jb

Stubbendieck, R. M., & Straight, P. D. (2015). Escape from Lethal Bacterial Competition through Coupled Activation of Antibiotic Resistance and a Mobilized Subpopulation. PLOS Genetics, 11(12), e1005722. https://doi.org/10.1371/journal.pgen.1005722

Taş, N., de Jong, A. E., Li, Y., Trubl, G., Xue, Y., & Dove, N. C. (2021). Metagenomic tools in microbial ecology research. Current Opinion in Biotechnology, 67, 184–191. https://doi.org/10.1016/J.COPBIO.2021.01.019

Vaz Jauri, P., & Kinkel, L. L. (2014). Nutrient overlap, genetic relatedness and spatial origin influence interaction-mediated shifts in inhibitory phenotype among Streptomyces spp. FEMS Microbiology Ecology, 90(1), 264–275. https://doi.org/10.1111/1574-6941.12389

Yim, G., Huimi Wang, H., & Davies, J. (2006). The truth about antibiotics. International Journal of Medical Microbiology, 296(2–3), 163–170. https://doi.org/10.1016/J.IJMM.2006.01.039

Wandersman, C., & Delepelaire, P. (2004, September 13). Bacterial iron sources: From siderophores to hemophores. Annual Review of Microbiology, Vol. 58, pp. 611–647. https://doi.org/10.1146/annurev.micro.58.030603.123811

Wootton, J. T. (1994). The nature and consequences of indirect effects in ecological communities. Annual Review of Ecology and Systematics, 25, 443–466. https://doi.org/10.1146/annurev.ecolsys.25.1.443

Zapién-Campos, R., Olmedo-álvarez, G., & Santillán, M. (2015). Antagonistic interactions are sufficient to explain self-assemblage of bacterial communities in a homogeneous environment: A computational modeling approach. Frontiers in Microbiology, 6(MAY), 1–9. https://doi.org/10.3389/fmicb.2015.00489

