## Supplementary figures and images for "A three-strain synthetic community model whose rapid response to antagonism allows the study of higher-order dynamics and emergent properties in minutes"

### Supplementary Fig 1

Supplementary Figure 1

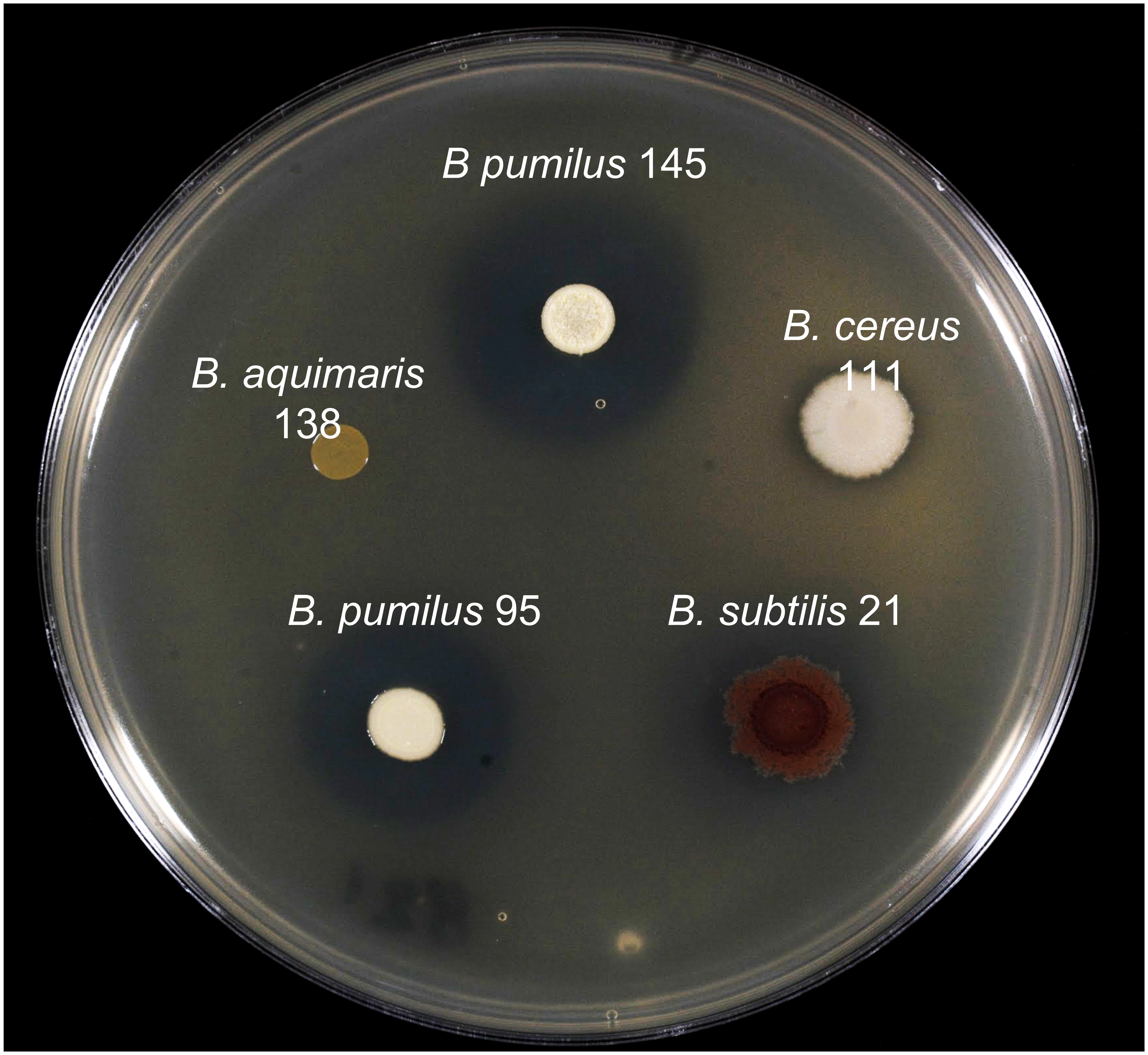

### Supplementary Fig 3

Supplementary Figure 3

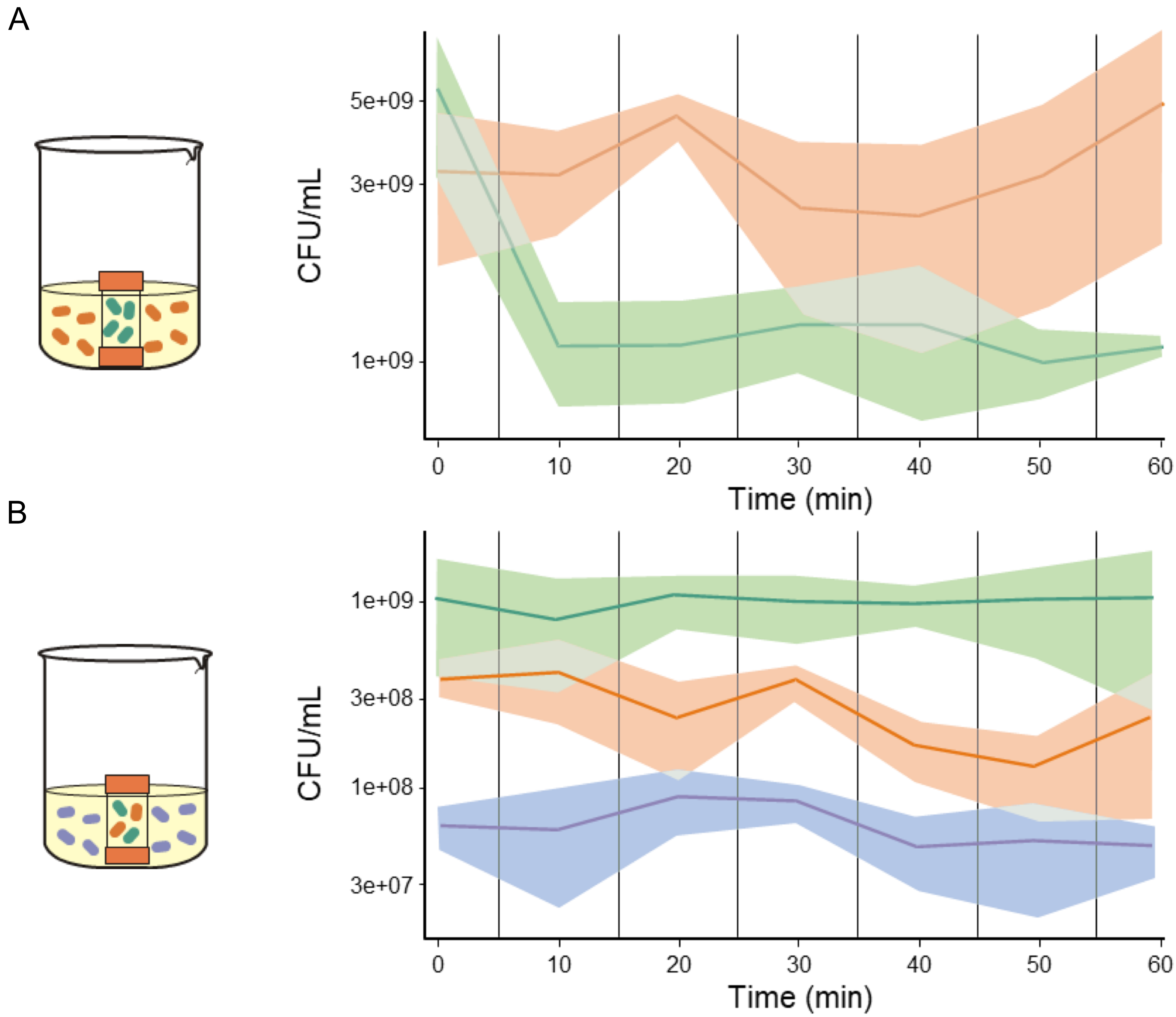

### Supplementary Fig 4

Supplementary Figure 4

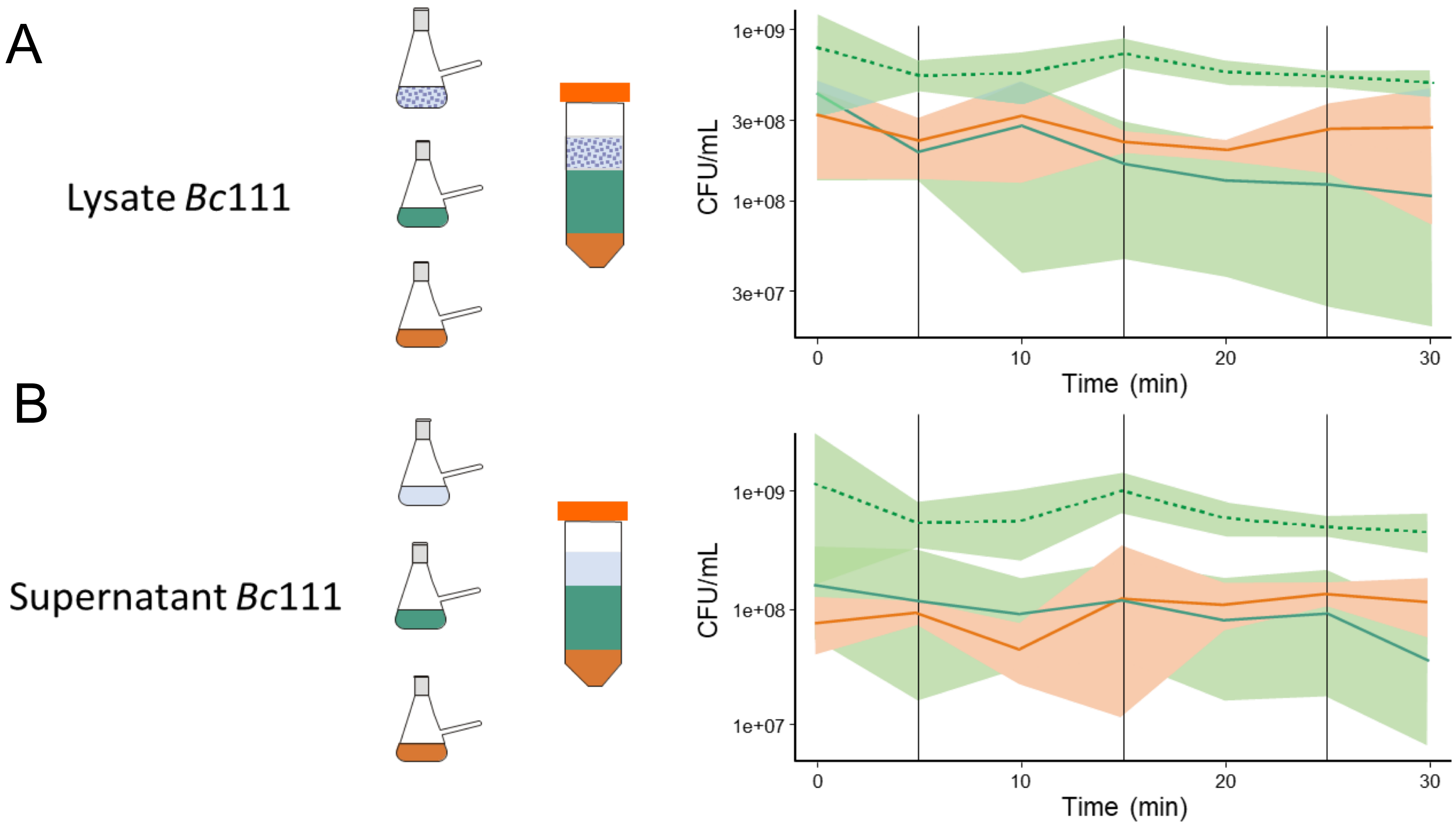
