## Supplementary Fig 2 for "A three-strain synthetic community model whose rapid response to antagonism allows the study of higher-order dynamics and emergent properties in minutes"

### Supplementary Figure 2

**A**

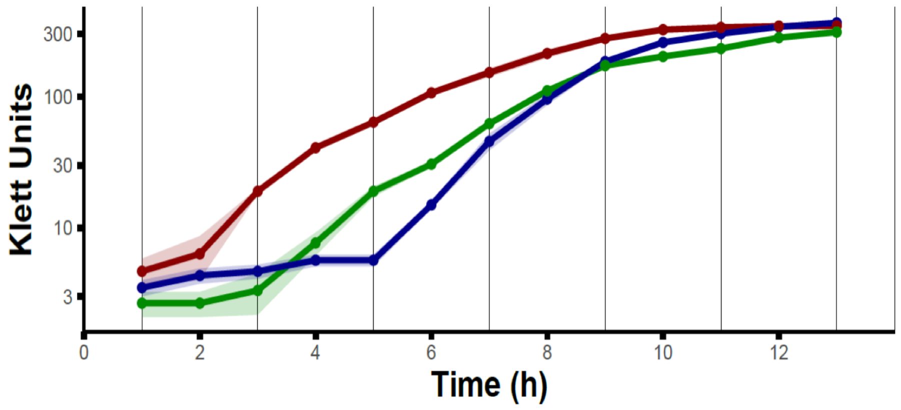


**B**

| **Strain** | **In-degree** | **Out-degree** | **Duplication rate** |
|---|---|---|---|
| *S. horikoshii* 20a | 18 | 0 | 38 min |
| *B. pumilus* 145 | 4 | 36 | 51 min |
| *B. cereus* 111 | 4 | 0 | 30 min |

**C**

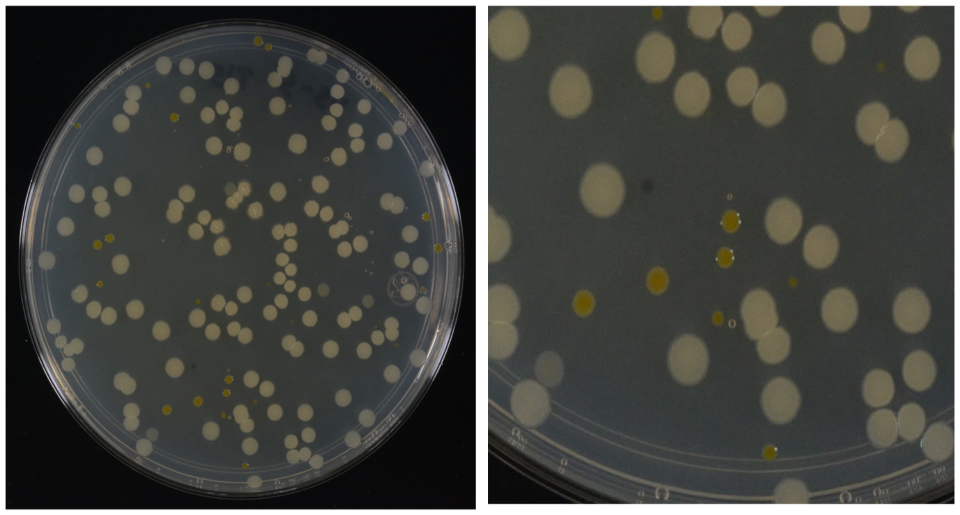

**D**

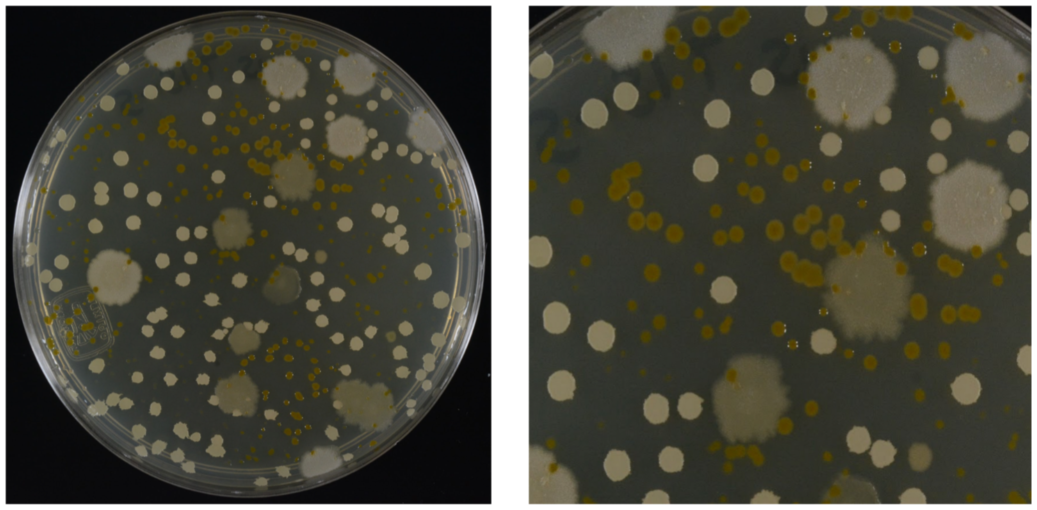
